# Cancer stem cell-derived extracellular vesicles preferentially target MHC-II– macrophages and PD1+ T cells in the tumor microenvironment

**DOI:** 10.1101/2022.04.26.489579

**Authors:** P. Gonzalez-Callejo, Z. Guo, T. Ziglari, N.M. Claudio, N. Oshimori, J. Seras-Franzoso, F. Pucci

## Abstract

Immunotherapy is an approved treatment option for head and neck squamous cell carcinoma (HNSCC). However, the response rate to immune checkpoint blockade is only 13% for recurrent HNSCC, highlighting the urgent need to better understand tumor-immune interplay, with the ultimate goal of improving patient outcomes. HNSCC present high local recurrence rates and therapy resistance that can be attributed to the presence of cancer stem cells (CSC) within tumors. CSC exhibit singular properties that enable them to avoid immune detection and eradication. The immune cell types that directly engage with CSC to allow immune escape and cancer recurrence are still unknown. Here, we genetically engineered CSC-derived extracellular vesicles (EVs) to perform sortase-mediated *in vivo* proximity labeling. We identified specific immune cell subsets recruited into the CSC niche. We demonstrated that unmanipulated CSC-EVs preferentially target MHC-II– macrophages and PD1+ T cells, and that such EV-mediated intercellular communication between CSC and these immune cells contributed to the observed spatial interactions and niche sharing. These results suggest that combination therapies targeting CSC, tumor macrophages and PD1 may synergize and lower local recurrence rates in HNSCC patients.

## Introduction

Head and neck squamous cell carcinoma (HNSCC) accounts for approximately 90% of oral and oropharyngeal cancer with over 400,000 new cases and more than 150,000 deaths reported each year worldwide.^1^ Advances in traditional treatments (surgery, radiotherapy, chemotherapy) have failed to increase survival due to patients presenting incurable advanced-stage disease and lymph node metastasis that ultimately cause their death.^2^

Immunotherapy is an approved treatment option for Head and neck squamous cell carcinoma (HNSCC).^3^ However, the response rate to immune checkpoint blockade is only 13% for recurrent HNSCC, highlighting the urgent need to better understand tumor-immune interplay, with the ultimate goal of improving patient outcomes.^3^ Click or tap here to enter text.

HNSCC present high local recurrence rates and therapy resistance that can be attributed to the presence of Cancer Stem Cells (CSC) within tumors. Several cell biomarkers such as CD44^4–6^, CD133^7,8^, SMAD Responsive Element (SRE)^9^and Aldehyde Dehydrogenase (ALDH) activity^10–13^ have identified specific CSC-like populations in HNSCC tumors with enhanced tumorigenic potential and resistance to chemo-or radiotherapy. CSC exhibit unique malignant intrinsic characteristics and play key roles in tumor initiation, growth and metastasis. CSC are also believed to drive therapy resistance and tumor relapse, as they can survive and dynamically adapt to changing and unfavorable environmental conditions.^14–19^

CSC exhibit singular properties that enable them to avoid immune detection and eradication.^20^ Recently, a number of studies have shown that CSC contribute to the generation of an immunosuppressive, pro-tumorigenic immune milieu by regulating the activity of various immune cells in the tumor microenvironment (TME). CSC can modulate T cells, tumor-associated macrophages (TAMs) and myeloid-derived suppressor cells activity towards immunosuppressive pathways.^20–25^ Importantly, these immune cells can also sustain CSC stemness and survival. ^25–28^ Such complex communication network between CSC and immune cells operates through various secreted cytokines, chemokines, growth factors and proteins of the extracellular matrix (ECM).^25,29^ Whether extracellular vesicles (EVs) also play a role is still unknown.

Emerging evidence has shown that tumors can interfere with host immunity by secreting EVs.^30^ EVs are defined as a heterogeneous collection of lipid bilayer membrane-enclosed vesicles naturally secreted by both prokaryotic and eukaryotic cells and that carry a complex cargo of mRNAs, lipids, metabolites, proteins and non-coding RNAs able to induce a response when signaling to EV-recipient cells.^31–39^ EVs of varying size, biogenesis and cargo content can be released from a single cell. Moreover, EV secretion pattern and content can change with changes in the physiological state of the parental cell.^40,41^ Once released, EVs can interact with cells in the immediate vicinity or at distant locations via transfer through lymphatic and blood circulation. Tumor derived EVs (tEVs) can affect the proliferation, apoptosis, cytokine production and reprogramming of both innate and adaptive immune cells, thereby modifying anti-cancer immune responses.^42–49^ Whether these functions belong to specific subpopulation of tEVs, such as those released by CSC, is still unclear.

## Materials and Methods

### Cell lines and culture conditions

#### Parental cell lines

Murine oral squamous cell carcinoma (OSCC) cell lines MOC2 (a chemical carcinogenesis model) and mEER (a Ras-dependent, HPV16-E6/E7-dependent model) were obtained from Kerafast, Inc, and Dr. Varner (UCSD), respectively. Both cell lines were routinely maintained in IMDM/DMEM/F12 (50:25:25) supplemented with 5% fetal bovine serum (FBS), 2 mM L-Glutamine, 1x Pen/Strep solution, Hydrocortizone (25ug/uL), Cholera Toxin (0.25ug/uL), Transferrin (25ug/uL), Insulin (10ug/uL), Tri-Iodo-Thyronine (0.2ug/uL), E.G.F. (10ug/mL). All cell cultures were propagated at 37°C and 5% CO_2_ in a humidified incubator.

#### Generation of modified cell lines expressing tEVs and tEVs ^CSC^ markers

mEER *dLNGFR:mCMV-PGK:CD63-eGFP* cell line was generated as previously reported by our group.^49^ Lentiviral transfer plasmids coding for ALDH1A1:CD63-eGFP, SRE:CD63-eGFP, ALDH1A1:SrtA and SS-mSca-LPETGG: *mCMV-PGK:CD63-eGFP* were designed in house, cloned by Genewiz and propagated in DH5a bacteria. Maxiprep was performed with Endo-free Macherey-Nagel kit. Unconcentrated lentiviral vectors were generated as previously reported by our group.^49^ MOC2 and mEER cells were seeded at a concentration of 10^5^ cells per well in a 6-well plate and transduced with lentiviral vector supernatants (1:1 ratio with complete media) in the presence of 1 µg/ml polybrene (Millipore).

#### Copy number assay

Total DNA was extracted from genetically modified 200,000 mEER and MOC2 *ALDH1A1:CD63-eGFP* and *SRE:CD63-eGFP* cell lines using QIAamp DNA Micro Kit (Qiagen). LV sequence was detected using a custom taqman assay (Applied Biosystems) on RRE sequence in a Viia7 PCR system. TaqMan probes for reference genes were ActinB, GusB and HPRT-1. One copy per genome standard was used, as previously described.^49^

#### Orosphere formation assay

5000 mEER and MOC cells/well were seeded in 6-well ultra-low attachment plates (Corner) in StemXVivo Serum-Free Tumorsphere Media (R&D Systems). Cells were cultured for 10 to 14 days, and orosphere formation was assessed in each well using light microscopy.

#### Stem gene profile validation

Total RNA was extracted from 300,000 mEER and MOC2 (flow-sorted as ALDH1A1:CD63-eGFP+/− and SRE:CD63-eGFP+/−cells) using the RNeasy Mini Kit (Qiagen) and the RNA obtained was reverse transcribed using a HighCapacity cDNA Reverse Transcription Kit (Thermo Fisher Scientific) according to the manufacturer⍰s instructions. The cDNA reverse transcription product was amplified with specific probes by qPCR using TaqMan method (Thermo Fisher Scientific). The reaction was performed in triplicate on a Viia7 Real time PCR system (Applied Biosystems). Relative normalized quantities (NRQ) of mRNA expression were calculated using the comparative Ct method (2^−ΔΔCt^) with two reference genes (GAPDH and Actin) used as endogenous controls with Excel software.

#### Mice and tumor challenge

Six-to eight-week-old B6 mice were purchased from Charles River Laboratories. For tumor challenge, parental and genetically modified mEER and MOC2 cells were intradermally injected (1×10^6^ in 50 µl of PBS) in the flank. After 10 days mice were euthanized and tumors collected for further analysis.

#### Tumors IF imaging

5 μm thick OCT microsections from experimental tumors were mounted on glass slides for immunofluorescent labeling. Briefly, after 15’ fixation with 4%PFA samples were washed in PBS-Tween 0.3% and primary antibodies, anti-rabbit eGFP (1:200, Abcam) anti-CD45 Biotin (1:200, Biolegend) and anti-F4/80 Alexafluor-647 (1:200, Biolegend) were supplemented in PBS/BSA 3 % (w/v) and incubated O.N. at 4°C. Samples were further washed 3 times in PBS-Tween 0.3% before the addition of secondary antibody. Goat Anti-rabbit AlexaFluor488 1:1000 and Streptavidin AlexaFluor568 1:500 were added and incubated 1h at RT. Slides were then washed and mounted with mounting media ProLong for visualization. Tumors were imaged using a Spinning Disk Confocal microscope (Yokogawa CSU-X1 on Zeiss Axio Observer).

#### Flow cytometry

Tumors were mechanically dissociated into single cell suspensions as previously described. ^49^ Cell suspensions were stained with conjugated antibodies (Biolegend, BD or eBiosciences) and Zombie aqua (Sigma). Following strategy was used to identify cells of interest:

- Tumor cells (CD63GFP+ Zombie aqua– CD45– CD31–)
- Endothelial cells (Zombie aqua– CD45– CD31+)
- B cells (Zombie aqua– CD45+ B220+)
- Macrophages MHC-II+(Zombie aqua– CD45+ CD11b+ F4/80+ II+)
- Macrophages MHC-II-(Zombie aqua– CD45+ CD11b+ F4/80+ II-)
- Inflammatory monocytes (Zombie aqua– CD45+ CD11b+ F4/80-CD11c+)
- Resident monocytes (Zombie aqua– CD45+ CD11b+ F4/80-CD11c-)
- Neutrophils (Zombie aqua– CD45+ F4/80-CD11c-SSChii)
- Dendritic cells (Zombie aqua– CD45+ CD11c+ F4/80– II+)
- PD-1 + T cells (Zombie aqua– CD45+ F4/80-B220-CD3+ PD-1+)
- PD-1 − T cells (Zombie aqua– CD45+ F4/80-B220-CD3+ PD-1-) Fluorochromes employed were the following: eGFP, Bv421, Bv605, Bv785, PE, PerCP, PC7, APC, A700, AC7.

#### Statistical analysis

Bar graphs display mean value ± standard error of the mean (SEM). 2-way ANOVA Holm-Sidak’s test or non-parametric Tukey’s test were employed for multiple mean comparisons. The significance threshold was established at p<0.05, and significance levels were schematically assigned *(0.01 ≤ p < 0.05), **(0.001 ≤ p < 0.01), ***(0.0001 ≤ p, ****(0.00001 ≤ p). All the analyses and graphs were performed using GraphPad Prism 6 software (GraphPad, San Diego).

## Results

### Genetic labeling of cancer stem cell-derived extracellular vesicles

Tumor secreted EVs (tEVs) represent prominent regulators of the immune response in cancer.^30^ CSC secreted EVs (tEVs^CSC^) are a subset of tEVs whose immunomodulating activity is still unknown. In order to start investigating whether tEVs^CSC^ have a role in shaping immune cell activity in the TME, we genetically labeled tEVs^CSC^ with fluorescent proteins. This approach allows to avoid any bias in EV composition due to *in vitro* isolation and assumptions on *in vivo* biodistribution of tEVs.^49^ In particular, we genetically engineered murine oral squamous cell carcinoma (OSCC) cell lines to express the vesicular membrane-associated protein CD63, fused with enhanced green fluorescence protein (CD63-eGFP) under the control of a CSC-specific promoter. We tested two different CSC-specific promoters, ALDH1A1 and SRE. ^9–13^ As reference controls, we genetically labeled the whole population of tEVs (including tEVs^CSC^) by expressing the CD63-eGFP fusion protein under a constitutive promoter (PGK). We worked on two different OSCC cell lines, a chemical carcinogenesis model (MOC2) and a Ras-dependent, HPV16-E6/E7-dependent model (mEER). MOC2 carry the same mutations observed in human HN cancers, namely Trp53, MAPK and FAT whereas mEER+ have been engineered to express Hras(G12) and HPV-E6/E7. Together, the mutational landscape of these two cell lines model >95% of human pathology. As expected, the constitutive reporter (*PGK:CD63-eGFP+)* showed green fluorescence in virtually all tumor cells **(Fig.1A)**. On the other hand, much less CD63-eGFP+ cells were observed in both MEER and MOC2 cells carrying the vectors *ALDH1A1:CD63-eGFP* **(Fig.1B; Fig.S1A)** and *SRE:CD63-eGFP* **(Fig.1C, Fig.S1B)**. In order to confirm that the observed differential expression of the tEVs reporter CD63-eGFP was due to the restricted expression of the *ALDH1A1* and SRE promoters among CSC (and not because of low transduction efficiency), a lentiviral vector (LV) copy number assay was performed. These analyses showed that mEER *SRE:CD63-eGFP* cells carried on average 30 LV copies per cell (CpC), and that 5 CpC were detected in mEER *ALDH1A1:CD63-eGFP* cells, indicating full transduction of the tumor cell populations **(Fig.1D**). Flow cytometry analysis revealed that, expectedly, positive control mEER *PGK:CD63-eGFP* cells showed high levels of eGFP fluorescence. eGFP fluorescence was also detected in lower levels in modified mEER *ALDH1A1:CD63-eGFP* and *SRE:CD63-eGFP* cells, suggesting that eGFP brightest cells may constitute the CSC population **(Fig.1E**). To test this hypothesis, we flow sorted the top 5% of the engineered cells based on eGFP intensity and evaluated their expression of stemness markers and ability to form orospheres in low-attachment culture. RT-qPCR assay revealed that both mEER *ALDH1A1:CD63-eGFP* bright and mEER *SRE:CD63-eGFP* bright cells showed significantly higher expression levels of the stemness markers ALDH1A1, Nanog and SOX2 when compared to eGFP-cells **(Fig.1F)**. Similarly, MOC2 *ALDH1A1:CD63-eGFP* bright and MOC2 *SRE:CD63-eGFP* bright cells showed significantly greater expression of the stemness marker *ALDH1A1* than eGFP-cells **(Fig.S1C)**. Importantly, flow sorted mEER *ALDH1A1:CD63-eGFP* bright cells efficiently formed orospheres when cultured in serum-free Low-Attachment (LA) conditions **(Fig 1. G)** while eGFP-cells were not able to form orospheres but showed small cellular aggregations (data not shown). Similarly, MOC2 *ALDH1A1:CD63-eGFP* bright cells formed bigger cell clusters than eGFP-cells when cultured in LA conditions **(Fig.S1D)**. To confirm that the orospheres we obtained in culture contain *bona fide* CSC, we performed gene expression analysis on mEER *ALDH1A1:CD63-eGFP* bright orospheres and found that they expressed significant higher levels of the stemness markers *ALDH1A1, Nanog, Oct-4, CD-133 and Sox-2* than unsorted mEER *ALDH1A1:CD63-eGFP* cells cultured in attachment conditions **(Fig 1. H)**. Altogether, these data confirm the ability of *ALDH1A1:CD63-eGFP* and *SRE:CD63-eGFP* expression cassettes to restrict expression of the tEV reporter within the CSC-enriched subpopulation of mEER and MOC2 cancer cell lines.

**Figure 1.**
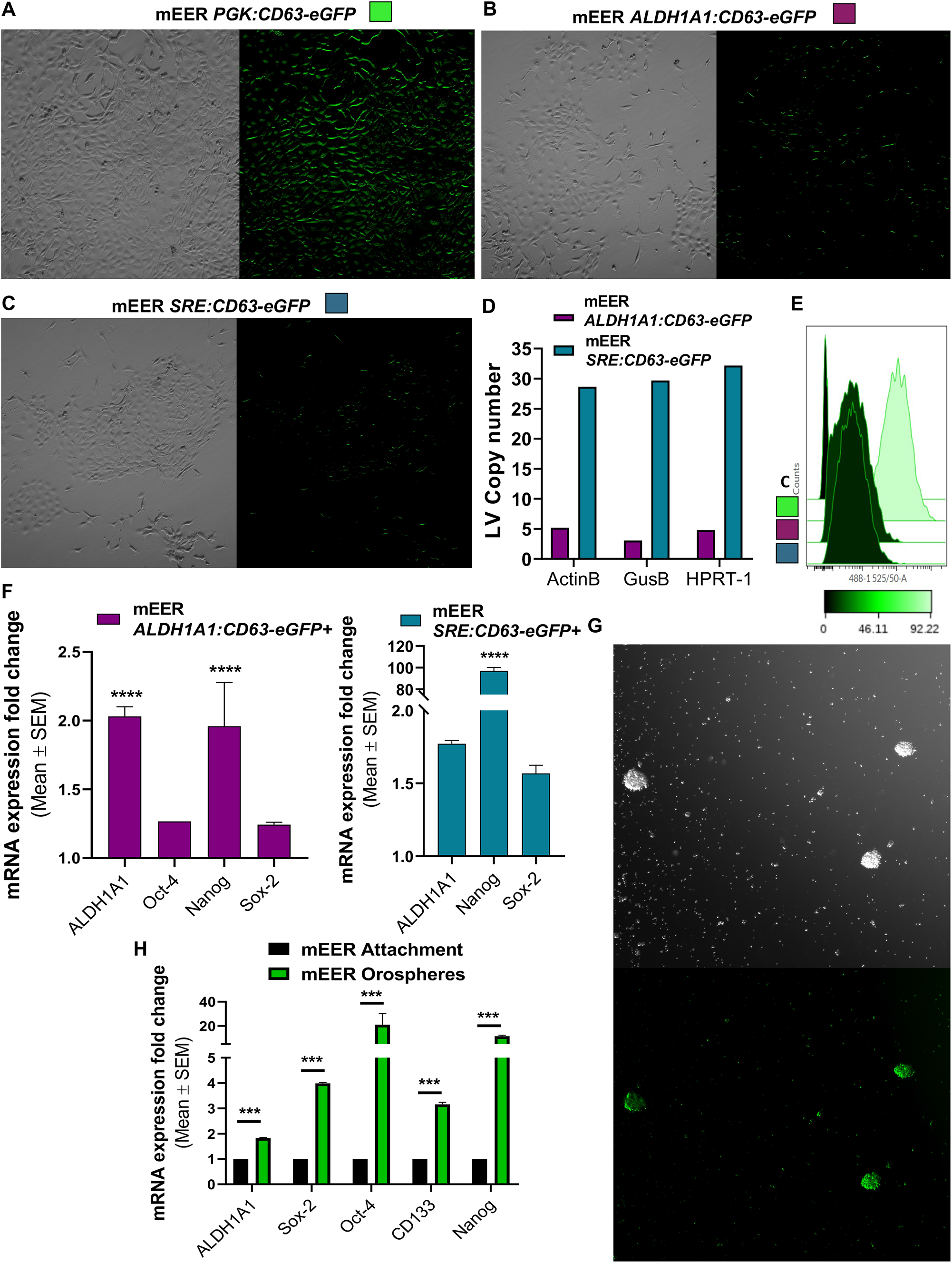
OSCC mEER CSC model characterization. Representative fluorescence confocal microscopy images of cultured mEER cells stably transduced with *PGK:CD63-eGFP* reporter **(A)**. Representative fluorescence confocal microscopy images of cultured mEER cells carrying the *ALDH1A1:CD63-eGFP* or the *SRE:CD63-eGFP* expression cassettes **(B, C)**. LV copy number present in genetically modified *ALDH1A1:CD63-eGFP* and *SRE:CD63-eGFP* mEER cells assessed by PCR **(D)**. eGFP fluorescence in unmodified **(C)** and genetically modified *PGK:CD63-eGFP* (green), *ALDH1A1:CD63-eGFP* (purple) and *SRE:CD63-eGFP* (teal) mEER cells analyzed by flow cytometry **(E)**. Relative increase in stemness gene expression of flow sorted brightest mEER eGFP + cells compared to eGFP-cells analyzed by RT-qPCR **(F)**. Representative microscopy images of orospheres growing from flow sorted *ALDH1A1:CD63-eGFP*+ mEER cultured in 3D tumorsphere-specific medium **(G)**. Stemness gene expression signature of mEER orospheres compared to mEER cells growing in attachment conditions assessed by RT-qPCR **(H)**. Holm-Sidak’s t test was used for statistical analysis.

### tEVs^csc^ preferentially target MHC-II– Macrophages and PD-1+ T cells

We and others have previously investigated the interactions that occur in the TME between tEVs and immune cells.^48,49^ Mononuclear phagocytes and tumor endothelial cells were among the cell types that bound tEVs at the highest rate. Whether tEVs^CSC^ possess a distinct tropism toward tumor infiltrating immune cells is still unknown. In order to test if tEVs^CSC^ preferentially interact with specific immune cell subsets, we challenged mice with mEER tumor cells carrying the *ALDH1A1:CD63-eGFP* or the *SRE:CD63-eGFP* expression cassette. As control, we used mice bearing mEER tumor cells carrying the *PGK:CD63-eGFP* expression cassette. Flow cytometry-based analysis of tumors revealed the presence of different levels of CD63-eGFP+ cells among groups **(Fig.2A)**. We then asked if differences in tEVs and tEVs^CSC^ tropism exist within functional subsets of CD45+ cells. Specifically, F4/80+ MHCII+ and F4/80+ MHCII-Macrophages (Mac), inflammatory and resident monocytes (Mo), PD-1+ and PD-1-T cells, Neutrophils (Neu), B cells and dendritic cells (DC) immune subpopulations were analyzed **(Fig.S2)**. In tumors formed by mEER cells constitutively expressing CD63-eGFP, CD45+ CD63-eGFP+ cells were composed mainly of MHC-II+ Mac (30.6%), B cells (16.5%) and Inflammatory Mo (16%), followed by Neu (12.9%) **(Fig.2B)**. When we analyzed tumors expressing either of the tEVs^CSC^ reporters, we observed an increased fraction of CD63-eGFP+ MHC-II– Mac among CD45+ CD63-eGFP+ cells in tEVs^CSC^ reporter tumors (27.8%, average between *ALDH1A1:CD63-eGFP and SRE:CD63-eGFP*) compared to *PGK:CD63-eGFP* tumors (5.7%). Interestingly, we observed an enrichment in the interactions between mEER tEVs^CSC^ and PD-1+ T cells (11.4%), as compared to tEVs (3.4%). **(Fig.2C)**. We observed statistically significant differences between the percentage of CD63-eGFP+ MHC-II+ Mac infiltrating tumors constitutively expressing the tEV reporter (87.4%) and those present in tumors carrying CSC reporters (10.5%), indicating that MHC-II+ Mac predominantly uptake non-CSC tEVs. On the other hand, the percentage of CD63-eGFP+ MHC-II– Mac did not significantly change, suggesting that MHC-II– Mac predominantly uptake tEVs^CSC^ **(Fig.2D)**. When we analyzed monocyte subsets, we observed a significant decrease in the percentage of CD63-eGFP+ monocytes from both inflammatory and resident subsets, indicating that monocytes predominantly uptake non-CSC tEVs. Similar results were observed with Neu, B cells and DC subsets, indicating that those populations preferably uptake non-CSC tEVs **(Fig.2D)**. By labeling T cells with the activation marker PD-1, we observed that the percentage of CD63-eGFP+ PD-1+ T cells did not significantly change between tumors constitutively expressing the tEVs reporter (9.6%) and those present in tumors carrying CSC reporters (6%), suggesting that, similarly to MHC-II– Mac, also PD-1+ T cells predominantly uptake tEVs^CSC^ **(Fig.2C)**. To highlight these differences, we calculated a tEVs^CSC^ specificity index by dividing the percentage of CD45+ CD63-eGFP+ immune cells for each subset in the tEVs^CSC^ groups by the percentage of the corresponding CD45+ CD63-eGFP+ subsets from the tEVs group. We observed that the index for MHC-II-Mac and PD-1+ T cells was significantly increased for those populations when compared to the index of all the other immune cell subsets. While tEVs^CSC^ specificity index mean was 0.87 for MHC-II– Mac and 0.63 for PD-1+ T cells, all other tested subsets were below 0.21 **(Fig2.E)**. Altogether, these data indicate that tEVs^CSC^ possess a preferential tropism towards MHC-II– Mac and PD-1+ T cells.

**Figure 2.**
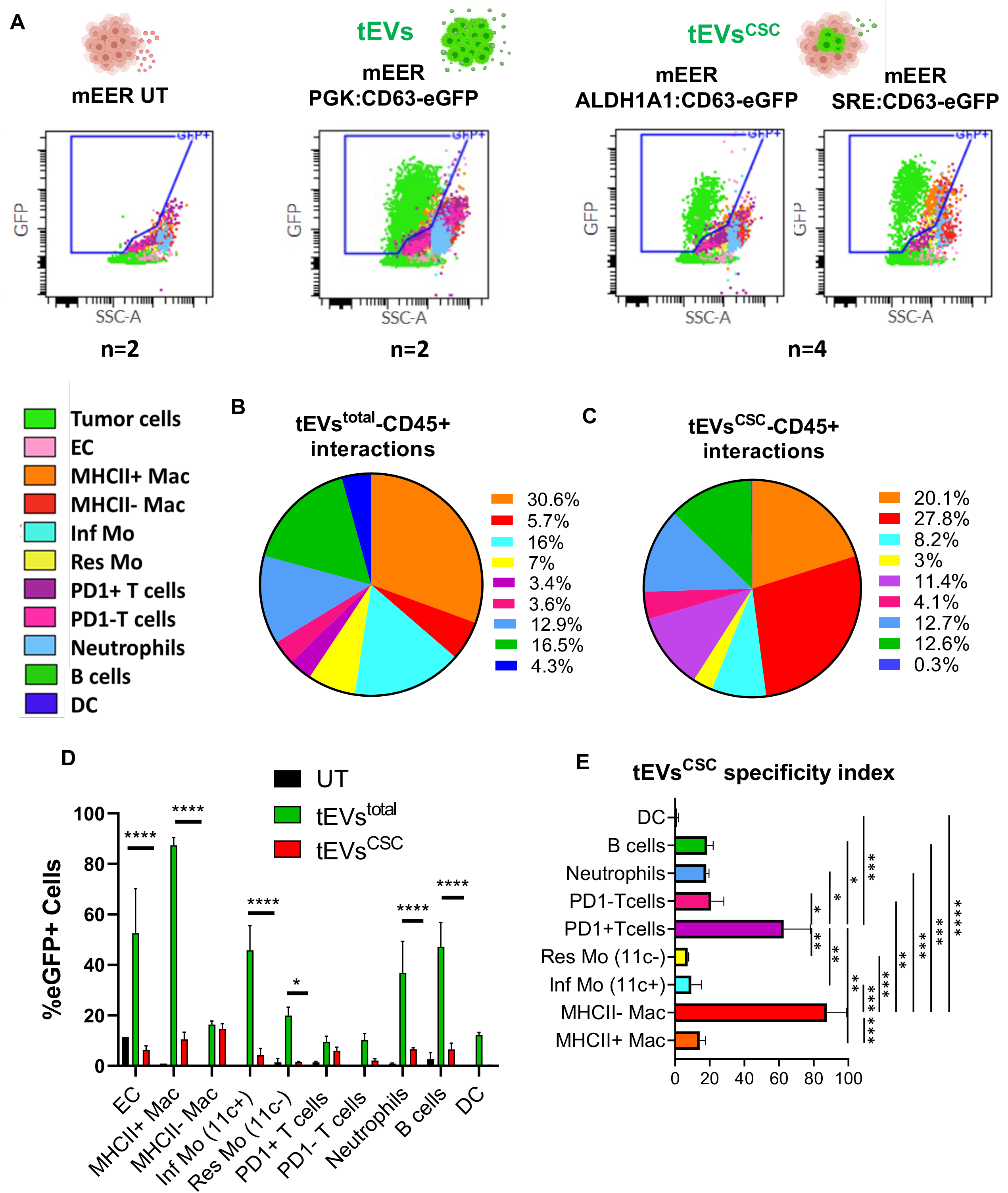
*in vivo* released mEER tEVs ^CSC^ target MHC-II-Mac and PD-1+ T cells in the TME. Representative overlaid plot of color-coded cell subsets present in tumors from mice bearing genetically modified mEER cells studied by flow cytometry. Sample number (n) for each group is indicated below each plot. CD63-eGFP+ gate is indicated in each case **(A)**. Representative graph of the % of specific tEVs–CD45+ immune cell subsets interactions **(B)** and for the % of specific tEVs^CSC^–CD45+ interactions **(C)**. Summary graph showing the % of cell subsets presenting CD63-eGFP+ fluorescence in unlabeled EVs tumors (UT), tEV-labeled tumors and tEV^CSC^-labeled tumors **(D)**. tEV^CSC^ specificity index showing the preferential interaction with tEVs^CSC^ by specific immune cell subsets **(E**). Two-way ANOVA Tukey’s and Holm-Sidak’s multiple comparisons tests were used for statistical analysis.

### Cancer stem cells and macrophages share the same niche within the tumor microenvironment

Our results so far indicate that MHC-II– Mac are selectively binding tEVs^CSC^. We next aimed to identify the mechanisms of such preferential binding. We hypothesized that CSC and MHC-II– Mac may share the same niches within the tumor microenvironment, which would increase exposure to tEVs^CSC^. We tested this hypothesis by imaging tumor sections. To this end, we performed immunofluorescence staining for CD45 and F4/80 on tumor sections from mice carrying either *PGK:CD63-eGFP* or *ALDH1A1:CD63-eGFP* expressing tumors. As expected, confocal microscopy images showed a broad CD63-eGFP+ signal in *PGK:CD63-eGFP* tumors **(Fig.3A)**, whereas fewer CD63-eGFP+ cells were found in *ALDH1A1:CD63-eGFP* tumors **(Fig.3B)**. Among CD45+ F480+ tumor macrophages, many were found at the tumor periphery. When considering those infiltrating the tumor mass, we observed a significant association of CD45+ F480+ tumor Mac with CSC, as quantified by measuring the distance between them in three distinct areas **(Fig.3C)**. These results suggest that the location of tumor macrophages may favor their preferential uptake of tEVs^CSC^.

**Figure 3.**
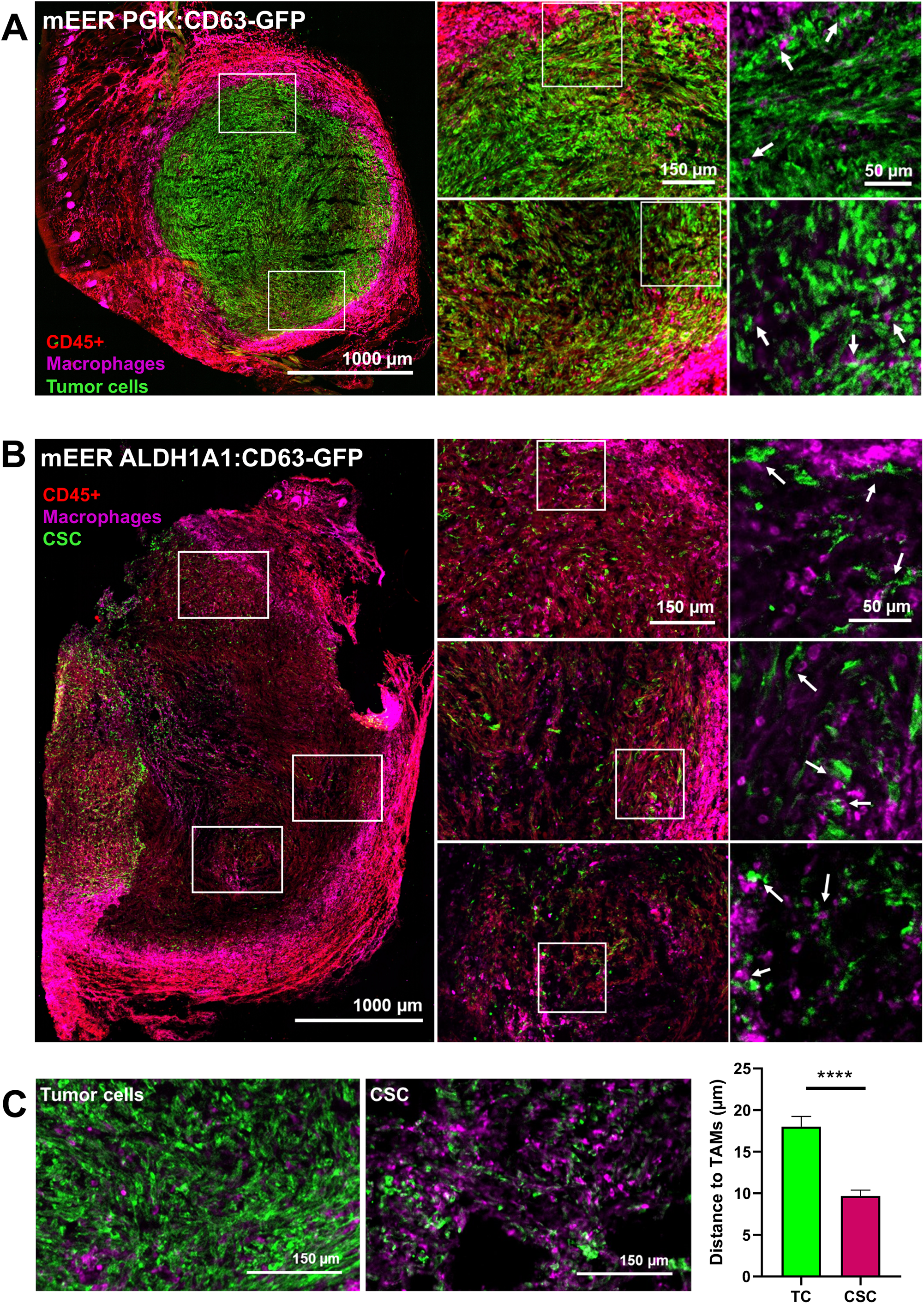
CSC show close localization to TAMs in the TME. Representative IF images of tumor sections carrying mEER *PGK:CD63-eGFP* tumor cells. Arrows in the insets indicate TAMs-tumor cells interactions (A). Representative IF images of tumor sections presenting mEER *ALDH1A1:CD63-eGFP* tumor cells. Arrows in the insets indicate TAMs-CSC interactions **(B)**. Examples of analyzed tumor areas employed to measure the distance in µm observed between tumor eGFP+ cells and TAMs using ImageJ software. Mann-Whitney test was used for statistical analysis **(C)**.

### Location-dependent labeling uncovers short-range interactions between CSC, tEVs^csc^ and MHC-II– Mac, PD-1+ T cells in the TME

To further investigate the ability of the CSC niche in shaping local immune cells, we employed a location-dependent labeling strategy we recently validated.^50^ The approach takes advantage of a membrane-bound bacterial transpeptidase, Sortase A (SrtA), expressed under the regulation of the ALDH1A1 promoter, to catalyze the transfer of a reporter on nearby immune cells **(Fig 4Ai)**. The reporter is monomeric Scarlett fluorescent protein (mSca) fused with SrtA recognition sequence (LPETGG) and with a secretory signal sequence, and is ubiquitously expressed by all tumor cells under the control of a constitutive bi-directional promoter, along with CD63-eGFP.^51^ The pan-tumor expression of CD63-eGFP served as internal control for total tumor-immune interactions. Since mSca is secreted in the extracellular environment, labeling is limited to near-by cells. Thus, this strategy allows to label host cells based on their proximity to CSC and their EVs. A detailed scheme of the experimental design is presented in **Fig.4Aii**. Flow cytometric analysis revealed that, as expected, the immune cell subsets most frequently interacting with tumor cells (as measured by tEV uptake) were MHC-II+ Mac (21.3% of total CD45+ CD63-GFP+ cells), followed by B cells (20.6%) and Neu (13%) **(Fig.4C)**, similarly to what we observed before **(Fig.2B)**. When the same tumors were analyzed to identify immune cell subsets found within the CSC niche, we found that the highest fraction of CD45+ mSca+ cells corresponded to MHC-II-Mac (30.5%), followed by MHC-II+ Mac (21.4%) and PD-1+ T cells (13%) **(Fig.4D)**. When we analyzed tumor-infiltrating CD45+ cell subsets presenting eGFP and mSca fluorescence, we observed a significant lower percentage of mSca+ cells compared to eGFP+ cells in all immune subsets indicating that they predominantly uptake non-CSC tEVs. Interestingly, we observed that the percentage of CD63-eGFP+ and mSca+ cells did not significantly change in MHC-II-Mac and PD-1+ T cells, suggesting that, similarly to previous results **(Fig. 2D)**, those subsets predominantly uptake tEVs^CSC^ **(Fig.4E)**. We then calculated a CSC niche-specificity index to summarize these results in one value. This index specifically increased for MHC-II-Mac (0.63) and PD-1+ T cell subpopulation (0.44), while it remained lower than 0.23 for the rest of the immune cell subsets **(Fig.4F)**. These results indicate that MHCII-Mac and PD1+ T cells dwell in proximity of CSC niches, which may explain their preferential uptake of tEVs^CSC^.

**Figure 4.**
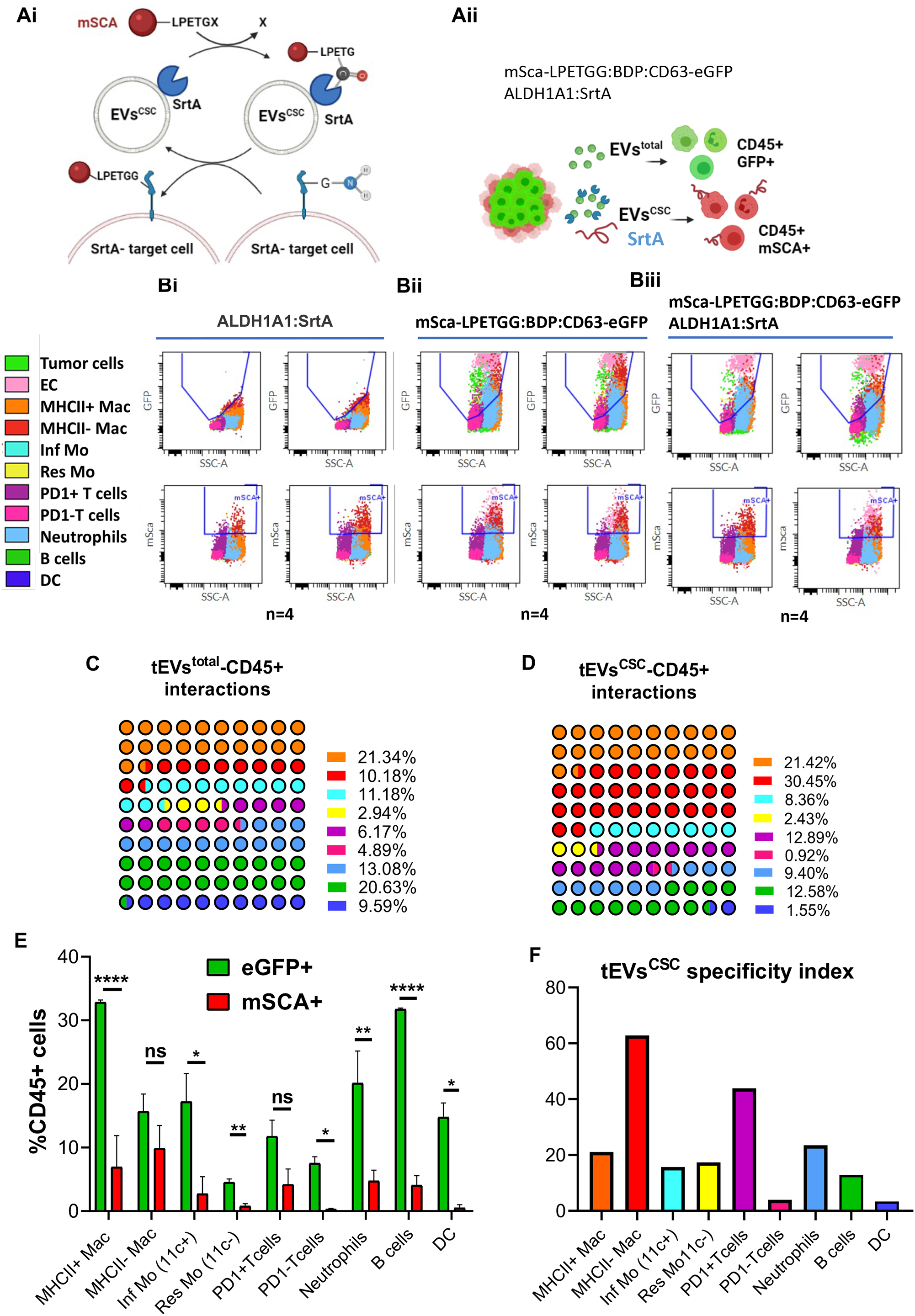
Second-degree labeling via tEVs^CSC^ reveals short range interactions between tEVs and MHC-II– Mac, PD-1+ T cells in the CSC niche. Illustrative schemes of SrtA enzymatic activity **(Ai)** and of the *in vivo* experimental design **(Aii)**. Overlaid plot representation of different cell populations present in tumors from mice bearing genetically modified mEER cells analyzed by flow cytometry. Two plots from n=4 are presented for each group. **(B)**. Representative graphs showing the % of eGFP+ immune cells in analyzed tumors for each subset with respect to all eGFP+ CD45+ cells **(C)**. Representative graphs showing the % of mSca+ immune cells for each subset with respect to all mSca+ CD45+ cells **(D)**. Summary graph presenting the % of immune cell subsets presenting eGFP+ and mSca+ fluorescence in tumors carrying modified mEER *ALDH1A1:SrtA/mSca-LPETGG:BDP:CD63-eGFP* tumor cells test. **(E)**. Calculated CSC niche-specificity index **(F)**. Two-way ANOVA Holm-Sidak’s multiple comparison test and multiple t tests were used for statistical analysis. BDP: Bi-directional promoter.

## Discussion

Tumor released EVs are key modulators of tumor immunity.^30^ The population of cancer cells is very heterogeneous, with some clones retaining stronger stem-like activity.^52,53^ Distinguishing between the contribution of CSC from that of more differentiated cancer cells to the total tEV pool has been very challenging, especially *in vivo*. A recent study used first-degree genetic labeling of tEV-targeted cells and focused on tumor cell-to-tumor cell signaling via EVs.^54^ The Authors uncovered an effective cooperation network mediated by tEVs and led by CSC, suggesting that a similar network may be in place between CSC and immune cells. Given the importance of CSC in cancer biology^19^, new technologies that allow to dissect the influence of native tEVs^CSC^ are necessary. In this study, we present a novel strategy to effectively track CSC secreted EVs in the TME under physiological conditions, avoiding any EVs *in vitro* manipulation. Here, we demonstrate proof-of-concept studies using first- and second-degree labeling of tEV-targeted cells in the TME, with a specific focus on tumor immune cell populations. We observed a surprising selectivity of tEVs^CSC^ in targeting specific immune cell subsets, namely MHC-II– Mac and PD-1+ T cells.

Analysis of tumors with fluorescently labeled tEVs and tEVs^CSC^ showed that the fraction of immune cell subsets presenting CD63-eGFP fluorescence was higher in the tEVs labeled group than in the tEVs^CSC^ labeled group. These results are expected since tEVs^CSC^ represent a minor fraction of bulk total tEVs. Deeper analysis of CD63-eGFP+ immune cell subpopulations in tumors identified Mac as the immune cell type with the highest interaction rate with both tEVs and tEVs^CSC^. These results are also expected, as TAMs constitute the most abundant population of tumor-infiltrating immune cells in TME.^55^ TAMs are educated by environmental factors to exhibit a spectrum of polarization phenotypes usually associated with specific functional states. One key functional biomarker of TAM polarization is MHC-II.^56^ Among the roles ascribed to MHC-II– Mac within the TME are immunosuppression,^57,58^ lymph/angiogenesis^59^, ECM deposition^60,61^and metastasis.^62^ Here, we report a clear difference between the interaction rates of MHC-II+ and MHC-II-Mac populations with each EVs fraction. While tEVs mainly interacted with MHC-II+ Mac, tEVs^CSC^ showed a significant preference toward MHC-II-Mac, as quantified by the specific interaction index. Numerous studies have shown the ability of tEVs to polarize TAMs towards pro-tumorigenic MHC-II-phenotypes.^63–66^ In this respect, our data suggest that tEVs^CSC^ may be the main tEV subset responsible for the reported TAMs polarization.

Our findings provide a potential mechanistic explanation of the recently reported maintenance of CSC niche by MHC-II– Mac.^26,28, 67^ The CSC niche is particularly important to support CSC self-renewal, repopulation potential, and tumor initiation.^68^ CSC contribute to the creation of a niche by inducing Mac polarization towards an immunosuppressive phenotype (MHC-II–), which in turn promotes and supports CSC aggressiveness.^69–72^ The relevance of TAMs in CSC biology is reinforced by a growing list of TAM-derived factors, including IL-6, IL-8, and CXCL1, that have been implicated in the maintenance of CSC stemness in different types of cancer.^69–74^ Here, we report that tEVs^CSC^ preferentially target MHC-II– Mac. In addition, novel second-degree (that is, sortase-based) labeling approaches demonstrated that CSC and MHC-II– Mac share the same niche, further indicating a role for tEVs in the creation of the CSC niche. As the presence of MHC-II– Mac and CSC populations in human tumors has been correlated with a poor prognosis for many types of cancer^57,75,76^, deeper knowledge of this communication network will be important to identify novel therapeutic opportunities.

Together with MHC-II– Mac, PD-1+ T cells also displayed specific interaction rates with tEVs^CSC^, as compared to total tEVs. The PD-1/PD-L1 pathway is a key immunosuppressive mechanism with significant clinical implications in many solid cancer types, including HNSCC. ^77,78^ Tumor EVs can also present PD-L1 on their surface, playing critical immunosuppressive roles when binding to PD-1+ T cells.^79,80^ Specifically, circulating PD-L1^high^ exosomes in HNCC patients’ plasma – but not soluble PD-L1 levels, have been associated with disease progression.^81^ Our study reveals that tEVs^CSC^ specifically interacted with PD-1+ T cell subsets, which may suggest the presence of PD-L1 ligand on tEVs^CSC^. We also show that PD-1+ T cells and CSC share the same niche. Thus, it is conceivable that tEVs^CSC^ may be primarily responsible for competing with immune checkpoint inhibitors in the clinic. Future study will confirm whether PD-L1 is enriched in CSC-derived tEVs and will assess if targeting PD-L1 on tEVs^CSC^ represents a novel therapeutic option. Alternatively, identification of the specific immune-modulators present on tEVs^CSC^ is needed to specifically target their immunosuppressive signals.

In conclusion, the present work not only establishes a novel technological platform to study tEVs^CSC^ and their roles in the TME at the single cell level, but also identifies specific immune cell subsets contributing to CSC biology. A better understanding of these microanatomical cross-talks will lead to the development of new therapeutic approaches with minimal to no side effects. Further studies focused on the immunomodulation orchestrated within the CSC niche promise to improve current immunotherapy approaches.

## Supplementary material

**Figure S1.**
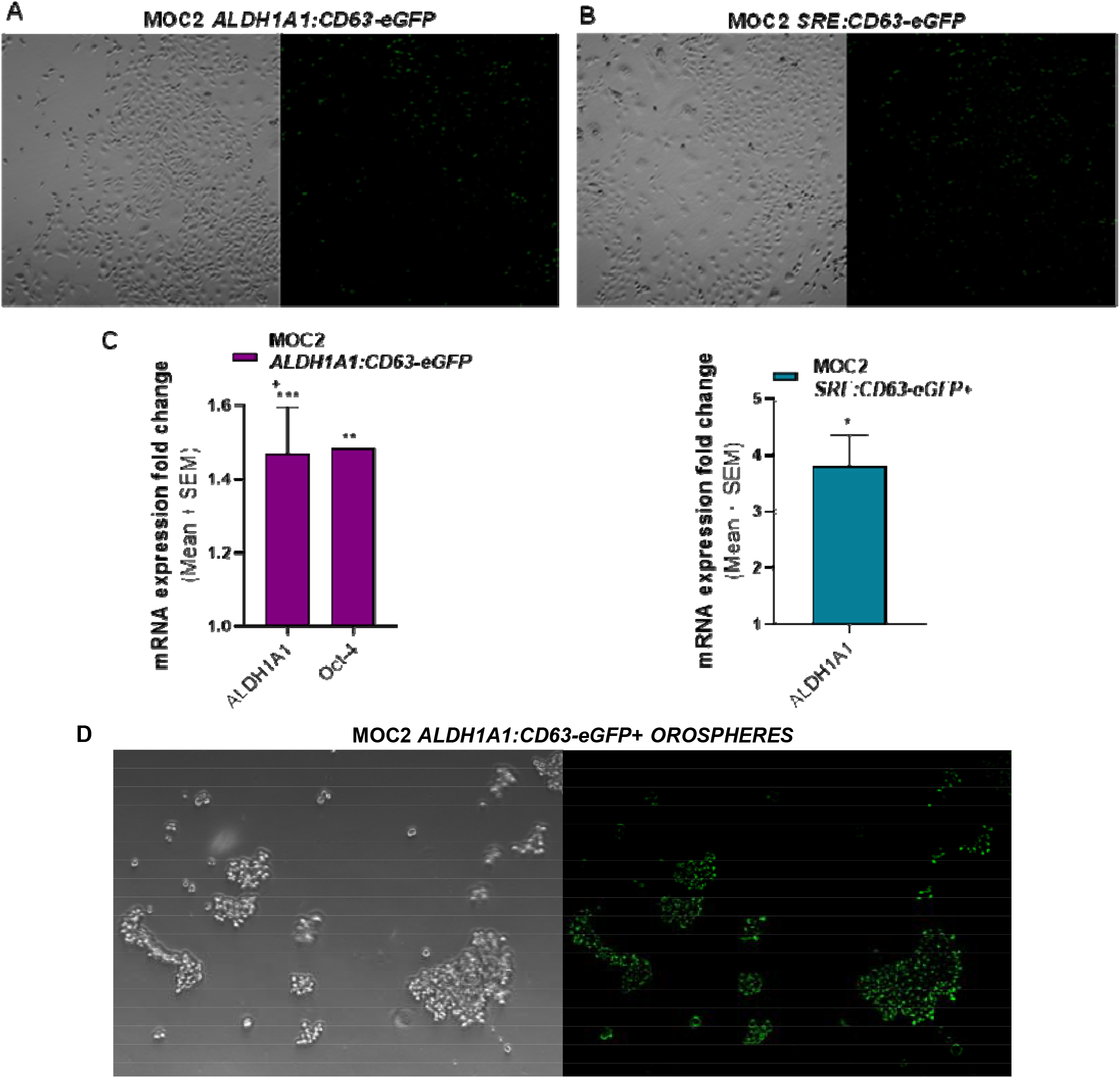
OSCC MOC2 CSC model. Representative confocal microscopy images of cultured *MOC2 ALDH1A1:CD63-eGFP* cells and *SRE:CD63-eGFP* cells in culture (A, B). Relative increase in stemness gene expression of flow sorted MOC2 eGFP + cells compared to eGFP-cells analyzed by RT-qPCR (C). Representative images of flow sorted MOC2 *ALDH1A1:CD63-eGFP+* cells growing in 3D tumorspheres specific medium **(D)**.

**Figure S2.**
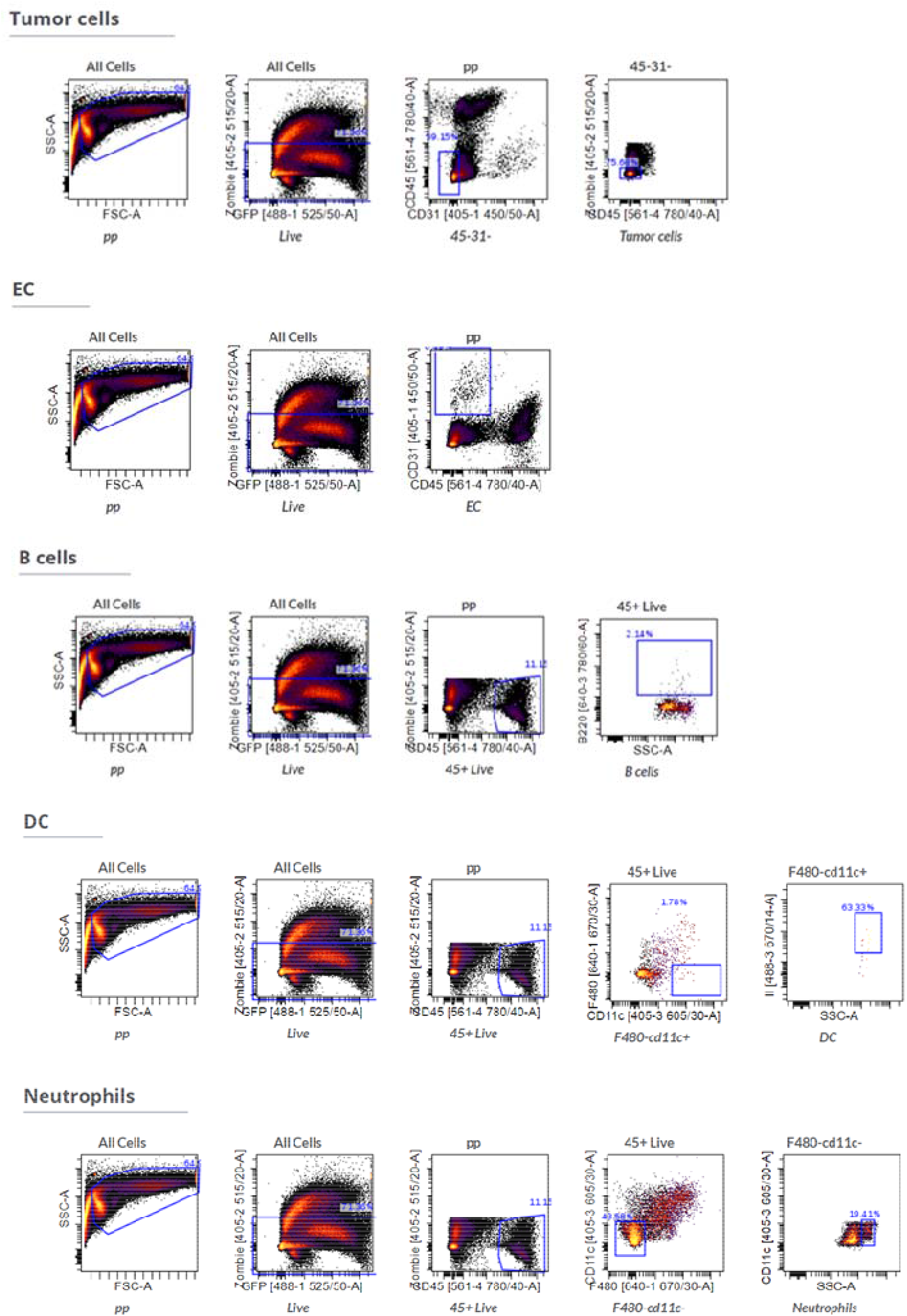
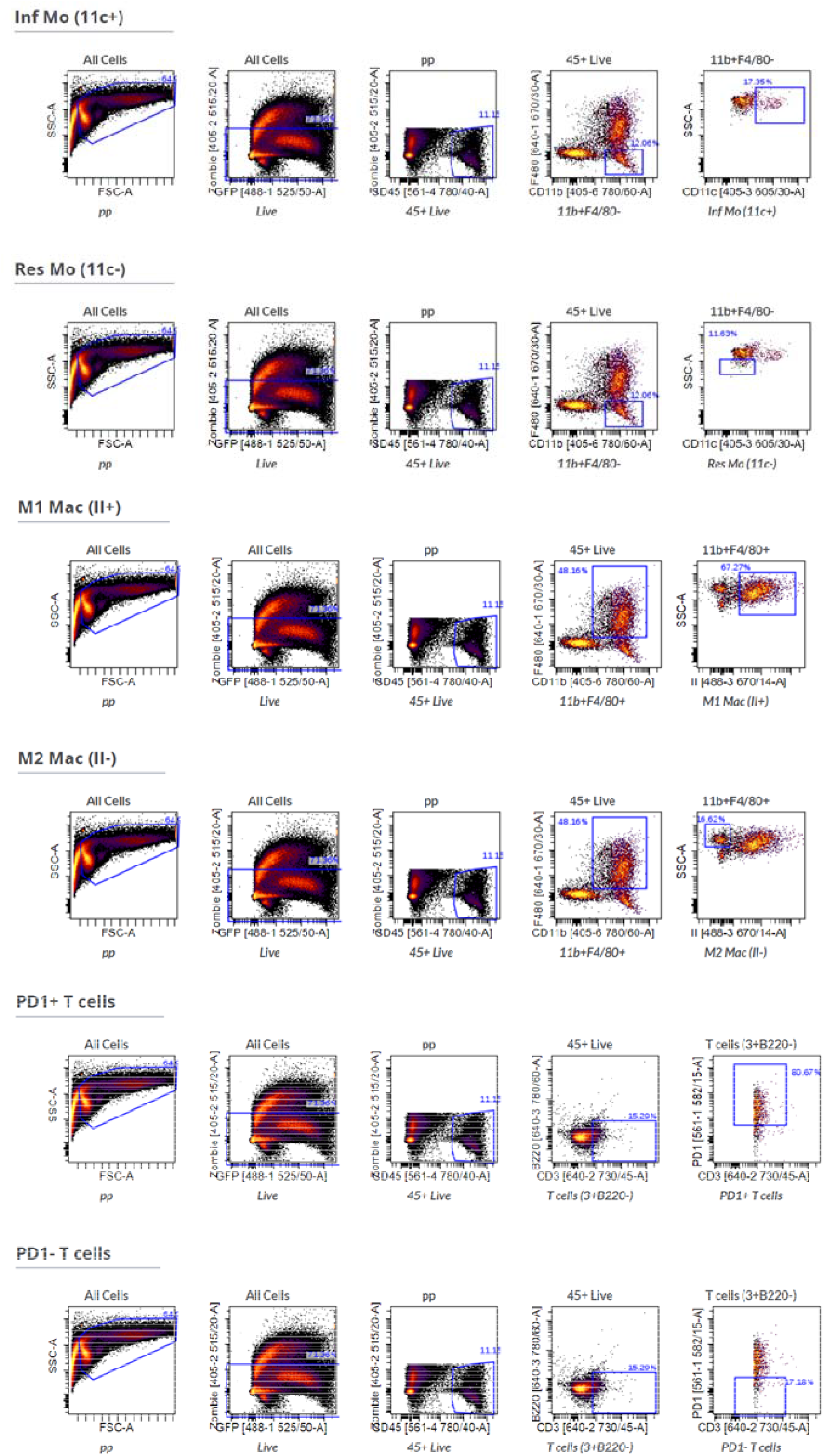
Flow cytometry gating analysis of studied tumors. Example of the gating strategy used to characterize the immune cells subsets present in analyzed tumors of *in vivo* experiments using Cytobank software.

**Figure S3.**
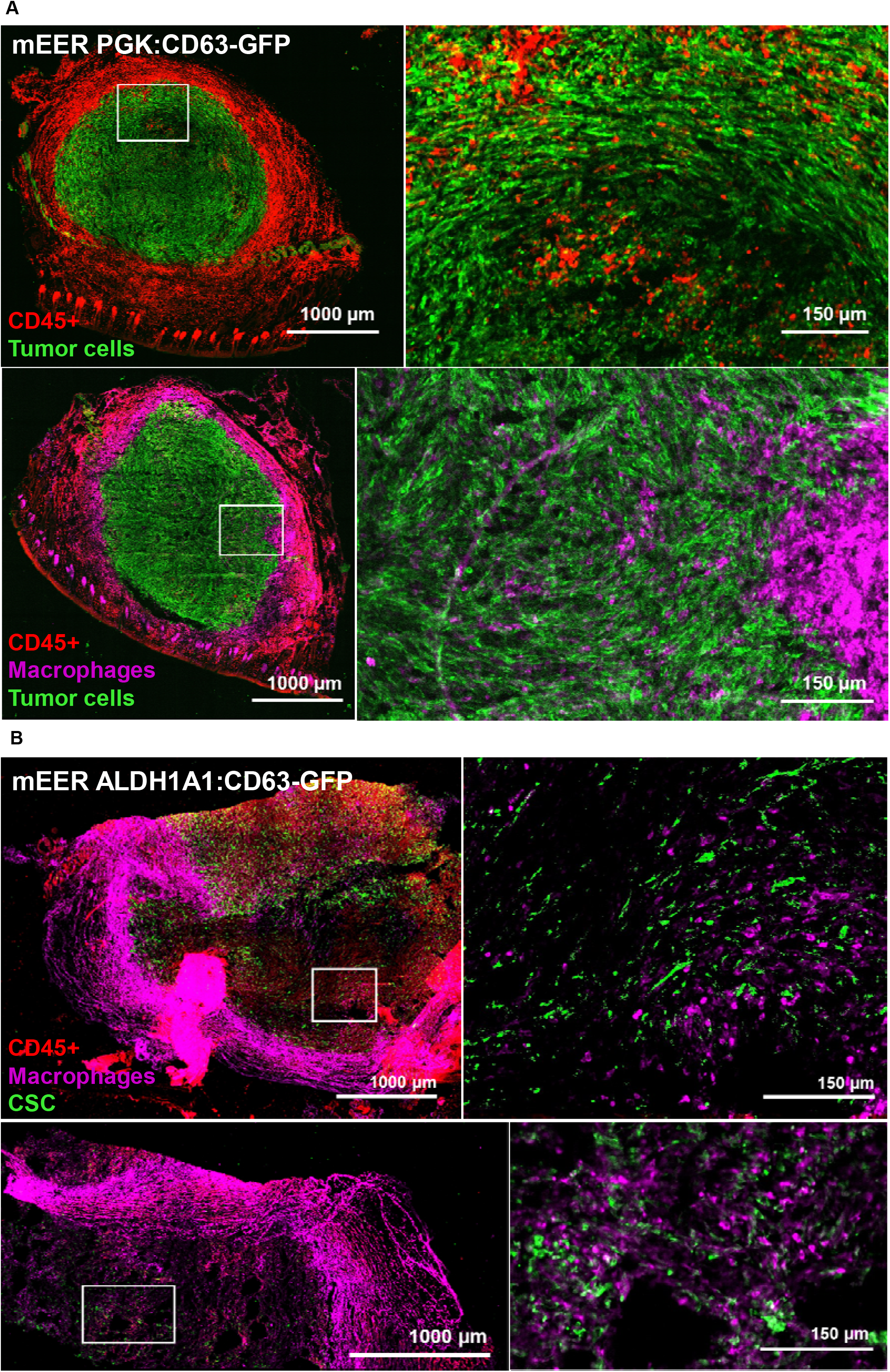
Additional representative IF images of tumor sections carrying mEER *CD63-eGFP+* and mEER *ALDH1A1:CD63-eGFP+* tumor cells.

## Bibliography

1. Sung, H. et al. Global Cancer Statistics 2020: GLOBOCAN Estimates of Incidence and Mortality Worldwide for 36 Cancers in 185 Countries. CA: A Cancer Journal for Clinicians (2021) doi:10.3322/caac.21660.

2. Johnson, D. E. et al. Head and neck squamous cell carcinoma. Nature Reviews Disease Primers (2020) doi:10.1038/s41572-020-00224-3.

3. Moskovitz, J., Moy, J. & Ferris, R. L. Immunotherapy for Head and Neck Squamous Cell Carcinoma. Current Oncology Reports vol. 20 (2018).

4. Prince, M. E. et al. Identification of a subpopulation of cells with cancer stem cell properties in head and neck squamous cell carcinoma. Proc Natl Acad Sci U S A (2007) doi:10.1073/pnas.0610117104.

5. Roy, S. et al. Inhibition of CD44 sensitizes cisplatin-resistance and affects Wnt/β-catenin signaling in HNSCC cells. International Journal of Biological Macromolecules (2020) doi:10.1016/j.ijbiomac.2020.01.131.

6. Gomez, K. E. et al. Cancer cell CD44 mediates macrophage/ monocyte-driven regulation of head and neck cancer stem cells. Cancer Research (2020) doi:10.1158/0008-5472.CAN-20-1079.

7. Wei, X. D. et al. In vivo investigation of CD133 as a putative marker of cancer stem cells in hep-2 cell line. Head and Neck (2009) doi:10.1002/hed.20935.

8. Wang, J. et al. Identification and characterization of CD133+CD44+ cancer stem cells from human laryngeal squamous cell carcinoma cell lines. J Cancer (2017) doi:10.7150/jca.17444.

9. Oshimori, N., Oristian, D. & Fuchs, E. TGF-β Promotes Heterogeneity and Drug Resistance in Squamous Cell Carcinoma. Cell 160, 963–976 (2015).

10. Chen, Y. C. et al. Aldehyde dehydrogenase 1 is a putative marker for cancer stem cells in head and neck squamous cancer. Biochemical and Biophysical Research Communications (2009) doi:10.1016/j.bbrc.2009.05.048.

11. Chen, Y. W. et al. Cucurbitacin I suppressed stem-like property and enhanced radiation-induced apoptosis in head and neck squamous carcinoma-derived CD44 +ALDH1+ cells. Molecular Cancer Therapeutics (2010) doi:10.1158/1535-7163.MCT-10-0504.

12. Qian, X. et al. ALDH1-positive cancer stem-like cells are enriched in nodal metastases of oropharyngeal squamous cell carcinoma independent of HPV status. Oncology Reports (2013) doi:10.3892/or.2013.2340.

13. Clay, M. R. et al. Single-marker identification of head and neck squamous cell carcinoma cancer stem cells with aldehyde dehydrogenase. Head and Neck (2010) doi:10.1002/hed.21315.

14. Reya, T., Morrison, S. J., Clarke, M. F. & Weissman, I. L. Stem cells, cancer, and cancer stem cells. Nature (2001) doi:10.1038/35102167.

15. Capp, J. P. Cancer stem cells: From historical roots to a new perspective. Journal of Oncology vol. 2019 (2019).

16. Kreso, A. & Dick, J. E. Evolution of the cancer stem cell model. Cell Stem Cell (2014) doi:10.1016/j.stem.2014.02.006.

17. Saxena, K., Murali, R., Kumar Jolly, M. & Nair, R. Cancer Stem Cell plasticity-a deadly deal. (2019) doi:10.20944/preprints201912.0388.v1.

18. Gilormini, M. et al. Isolation and characterization of a head and neck squamous cell carcinoma subpopulation having stem cell characteristics. Journal of Visualized Experiments (2016) doi:10.3791/53958.

19. Batlle, E. & Clevers, H. Cancer stem cells revisited. Nature Publishing Group 23, (2017).

20. Müller, L. et al. Bidirectional Crosstalk Between Cancer Stem Cells and Immune Cell Subsets. Frontiers in Immunology (2020) doi:10.3389/fimmu.2020.00140.

21. Clara, J. A., Monge, C., Yang, Y. & Takebe, N. Targeting signalling pathways and the immune microenvironment of cancer stem cells — a clinical update. Nature Reviews Clinical Oncology (2020) doi:10.1038/s41571-019-0293-2.

22. Wang, G. et al. Tumor microenvironment in head and neck squamous cell carcinoma: Functions and regulatory mechanisms. Cancer Letters (2021) doi:10.1016/j.canlet.2021.03.009.

23. Lee, Y. et al. Cd44+ cells in head and neck squamous cell carcinoma suppress t-cell-mediated immunity by selective constitutive and inducible expression of PD-L1. Clinical Cancer Research (2016) doi:10.1158/1078-0432.CCR-15-2665.

24. Chikamatsu, K., Takahashi, G., Sakakura, K., Ferrone, S. & Masuyama, K. Immunoregulatory properties of CD44+ cancer stem-like cells in squamous cell carcinoma of the head and neck. Head and Neck (2011) doi:10.1002/hed.21420.

25. Chen, P., Hsu, W. H., Han, J., Xia, Y. & DePinho, R. A. Cancer Stemness Meets Immunity: From Mechanism to Therapy. Cell Reports (2021) doi:10.1016/j.celrep.2020.108597.

26. Li, X. et al. CXCL12/CXCR4 pathway orchestrates CSC-like properties by CAF recruited tumor associated macrophage in OSCC. Experimental Cell Research (2019) doi:10.1016/j.yexcr.2019.03.013.

27. Nusblat, L. M., Carroll, M. J. & Roth, C. M. Crosstalk between M2 macrophages and glioma stem cells. Cellular Oncology (2017) doi:10.1007/s13402-017-0337-5.

28. Tao, W. et al. Dual Role of WISP1 in maintaining glioma stem cells and tumor-supportive macrophages in glioblastoma. Nature Communications (2020) doi:10.1038/s41467-020-16827-z.

29. Celià-Terrassa, T. & Kang, Y. Metastatic niche functions and therapeutic opportunities. Nature Cell Biology vol. 20 868–877 (2018).

30. Marar, C., Starich, B. & Wirtz, D. Extracellular vesicles in immunomodulation and tumor progression. Nature Immunology (2021) doi:10.1038/s41590-021-00899-0.

31. Maas, S. L. N., Breakefield, X. O. & Weaver, A. M. Extracellular Vesicles: Unique Intercellular Delivery Vehicles. Trends in Cell Biology 27, 172–188 (2017).

32. Desrochers, L. M., Antonyak, M. A. & Cerione, R. A. Extracellular Vesicles: Satellites of Information Transfer in Cancer and Stem Cell Biology. Developmental Cell (2016) doi:10.1016/j.devcel.2016.04.019.

33. Yáñez-Mó, M. et al. Biological properties of extracellular vesicles and their physiological functions. Journal of Extracellular Vesicles (2015) doi:10.3402/jev.v4.27066.

34. Van Niel, G., D’Angelo, G. & Raposo, G. Shedding light on the cell biology of extracellular vesicles. Nature Reviews Molecular Cell Biology (2018) doi:10.1038/nrm.2017.125.

35. Iraci, N., Leonardi, T., Gessler, F., Vega, B. & Pluchino, S. Focus on extracellular vesicles: Physiological role and signalling properties of extracellular membrane vesicles. International Journal of Molecular Sciences (2016) doi:10.3390/ijms17020171.

36. György, B. et al. Membrane vesicles, current state-of-the-art: Emerging role of extracellular vesicles. Cellular and Molecular Life Sciences (2011) doi:10.1007/s00018-011-0689-3.

37. Willms, E. et al. Cells release subpopulations of exosomes with distinct molecular and biological properties. Scientific Reports (2016) doi:10.1038/srep22519.

38. Yoon, Y. J., Kim, O. Y. & Gho, Y. S. Extracellular vesicles as emerging intercellular communicasomes. BMB Reports (2014) doi:10.5483/BMBRep.2014.47.10.164.

39. Zaborowski, M. P., Balaj, L., Breakefield, X. O. & Lai, C. P. Extracellular Vesicles: Composition, Biological Relevance, and Methods of Study. BioScience (2015) doi:10.1093/biosci/biv084.

40. Wen, S. W. et al. Breast Cancer-Derived Exosomes Reflect the Cell-of-Origin Phenotype. Proteomics 19, (2019).

41. Abels, E. R. & Breakefield, X. O. Introduction to Extracellular Vesicles: Biogenesis, RNA Cargo Selection, Content, Release, and Uptake. Cellular and Molecular Neurobiology (2016) doi:10.1007/s10571-016-0366-z.

42. Zhou, X. et al. The function and clinical application of extracellular vesicles in innate immune regulation. Cellular and Molecular Immunology (2020) doi:10.1038/s41423-020-0391-1.

43. Srivastava, A., Rathore, S., Munshi, A. & Ramesh, R. Extracellular Vesicles in Oncology: from Immune Suppression to Immunotherapy. AAPS Journal (2021) doi:10.1208/s12248-021-00554-4.

44. Wieckowski, E. U. et al. Tumor-Derived Microvesicles Promote Regulatory T Cell Expansion and Induce Apoptosis in Tumor-Reactive Activated CD8 + T Lymphocytes. The Journal of Immunology (2009) doi:10.4049/jimmunol.0900970.

45. Chen, X. et al. Exosomes derived from hypoxic epithelial ovarian cancer cells deliver microRNAs to macrophages and elicit a tumor-promoted phenotype. Cancer Letters (2018) doi:10.1016/j.canlet.2018.08.001.

46. Cooks, T. et al. Mutant p53 cancers reprogram macrophages to tumor supporting macrophages via exosomal miR-1246. Nature Communications (2018) doi:10.1038/s41467-018-03224-w.

47. Mittal, S., Gupta, P., Chaluvally-Raghavan, P. & Pradeep, S. Emerging role of extracellular vesicles in immune regulation and cancer progression. Cancers (Basel) (2020) doi:10.3390/cancers12123563.

48. Raimondo, S., Pucci, M., Alessandro, R. & Fontana, S. Extracellular vesicles and tumor-immune escape: Biological functions and clinical perspectives. International Journal of Molecular Sciences 21, (2020).

49. Pucci, F. et al. SCS macrophages suppress melanoma by restricting tumor-derived vesicle-B cell interactions. Science (1979) (2016) doi:10.1126/science.aaf1328.

50. Hamilton, N., Claudio, N. M., Armstrong, R. J. & Pucci, F. Cell Surface Labeling by Engineered Extracellular Vesicles. Advanced Biosystems 4, (2020).

51. Amendola, M., Venneri, M. A., Biffi, A., Vigna, E. & Naldini, L. Coordinate dual-gene transgenesis by lentiviral vectors carrying synthetic bidirectional promoters. Nature Biotechnology 23, (2005).

52. Reya, T., Morrison, S. J., Clarke, M. F. & Weissman, I. L. Stem cells, cancer, and cancer stem cells. Nature (2001) doi:10.1038/35102167.

53. Visvader, J. E. & Lindeman, G. J. Cancer stem cells: Current status and evolving complexities. Cell Stem Cell 10, 717–728 (2012).

54. Ruivo, C. F. et al. Extracellular Vesicles from Pancreatic Cancer Stem Cells Lead an Intratumor Communication Network (EVNet) to fuel tumour progression. Gut gutjnl- 2021-324994 (2022) doi:10.1136/gutjnl-2021-324994.

55. Baig, M. S. et al. Tumor-derived exosomes in the regulation of macrophage polarization. Inflammation Research (2020) doi:10.1007/s00011-020-01318-0.

56. Boutilier, A. J. & Elsawa, S. F. Macrophage polarization states in the tumor microenvironment. International Journal of Molecular Sciences vol. 22 (2021).

57. Pollard, R. & W., J. Tumor-associated macrophages⍰: from mechanisms to therapy. Immunity. (2015) doi:10.1016/j.immuni.2014.06.010.Tumor-associated.

58. Colegio, O. R. et al. Functional polarization of tumour-associated macrophages by tumour-derived lactic acid. Nature (2014) doi:10.1038/nature13490.

59. Riabov, V. et al. Role of tumor associated macrophages in tumor angiogenesis and lymphangiogenesis. Frontiers in Physiology vol. 5 MAR (2014).

60. Deligne, C. & Midwood, K. S. Macrophages and Extracellular Matrix in Breast Cancer: Partners in Crime or Protective Allies? Frontiers in Oncology vol. 11 (2021).

61. Ahmad, R. S., Eubank, T. D., Lukomski, S. & Boone, B. A. Immune cell modulation of the extracellular matrix contributes to the pathogenesis of pancreatic cancer. Biomolecules vol. 11 (2021).

62. Lin, Y., Xu, J. & Lan, H. Tumor-associated macrophages in tumor metastasis: Biological roles and clinical therapeutic applications. Journal of Hematology and Oncology vol. 12 (2019).

63. Zhou, Z. et al. CCL18 secreted from M2 macrophages promotes migration and invasion via the PI3K/Akt pathway in gallbladder cancer. Cellular Oncology (2019) doi:10.1007/s13402-018-0410-8.

64. Cai, J., Qiao, B., Gao, N., Lin, N. & He, W. Oral squamous cell carcinoma-derived exosomes promote M2 subtype macrophage polarization mediated by exosome-enclosed miR-29a-3p. American Journal of Physiology - Cell Physiology (2019) doi:10.1152/ajpcell.00366.2018.

65. Li, L. et al. Exosomes derived from hypoxic oral squamous cell carcinoma cells deliver miR-21 to normoxic cells to elicit a prometastatic phenotype. Cancer Research (2016) doi:10.1158/0008-5472.CAN-15-1625.

66. Ito, A. et al. Extracellular vesicles shed from gastric cancer mediate protumor macrophage differentiation. BMC Cancer (2021) doi:10.1186/s12885-021-07816-6.

67. Taniguchi, S. et al. Tumor-initiating cells establish an IL-33–TGF-b niche signaling loop to promote cancer progression. Science (1979) 369, (2020).

68. Plaks, V., Kong, N. & Werb, Z. The cancer stem cell niche: How essential is the niche in regulating stemness of tumor cells? Cell Stem Cell vol. 16 225–238 (2015).

69. Kong, L. et al. Deletion of interleukin-6 in monocytes/macrophages suppresses the initiation of hepatocellular carcinoma in mice. Journal of Experimental and Clinical Cancer Research (2016) doi:10.1186/s13046-016-0412-1.

70. Xu, X., Ye, J., Huang, C., Yan, Y. & Li, J. M2 macrophage-derived IL6 mediates resistance of breast cancer cells to hedgehog inhibition. Toxicology and Applied Pharmacology (2019) doi:10.1016/j.taap.2018.12.013.

71. Yin, Y. et al. The immune-microenvironment confers chemoresistance of colorectal cancer through macrophage-derived IL6. Clinical Cancer Research (2017) doi:10.1158/1078-0432.CCR-17-1283.

72. Zhu, X. et al. IL-6R/STAT3/miR-204 feedback loop contributes to cisplatin resistance of epithelial ovarian cancer cells. Oncotarget (2017) doi:10.18632/oncotarget.16610.

73. Lu, H. et al. A breast cancer stem cell niche supported by juxtacrine signalling from monocytes and macrophages. Nature Cell Biology (2014) doi:10.1038/ncb3041.

74. Mitchem, J. B. et al. Targeting tumor-infiltrating macrophages decreases tumor-initiating cells, relieves immunosuppression, and improves chemotherapeutic responses. Cancer Research (2013) doi:10.1158/0008-5472.CAN-12-2731.

75. Park, I. H. et al. Tumor-derived induces expression on immunosuppressive NK cells in triple-negative breast cancer. 8, 32722–32730 (2017).

76. Franklin, R. A. et al. The cellular and molecular origin of tumor-associated macrophages. Science (1979) (2014) doi:10.1126/science.1252510.

77. Schneider, S. et al. PD-1 and PD-L1 expression in HNSCC primary cancer and related lymph node metastasis – impact on clinical outcome. Histopathology 73, (2018).

78. Bai, J. et al. Regulation of PD-1/PD-L1 pathway and resistance to PD-1/PDL1 blockade. Oncotarget vol. 8 (2017).

79. Poggio, M. et al. Suppression of Exosomal PD-L1 Induces Systemic Anti-tumor Immunity and Memory. Cell (2019) doi:10.1016/j.cell.2019.02.016.

80. Chen, G. et al. Exosomal PD-L1 contributes to immunosuppression and is associated with anti-PD-1 response. Nature (2018) doi:10.1038/s41586-018-0392-8.

81. Theodoraki, M. N., Yerneni, S. S., Hoffmann, T. K., Gooding, W. E. & Whiteside, T. L. Clinical significance of PD-L1 þ exosomes in plasma of head and neck cancer patients. Clinical Cancer Research (2018) doi:10.1158/1078-0432.CCR-17-2664.

