## Supplementary material for "Cancer stem cell-derived extracellular vesicles preferentially target MHC-II– macrophages and PD1+ T cells in the tumor microenvironment": Suppl. Figures

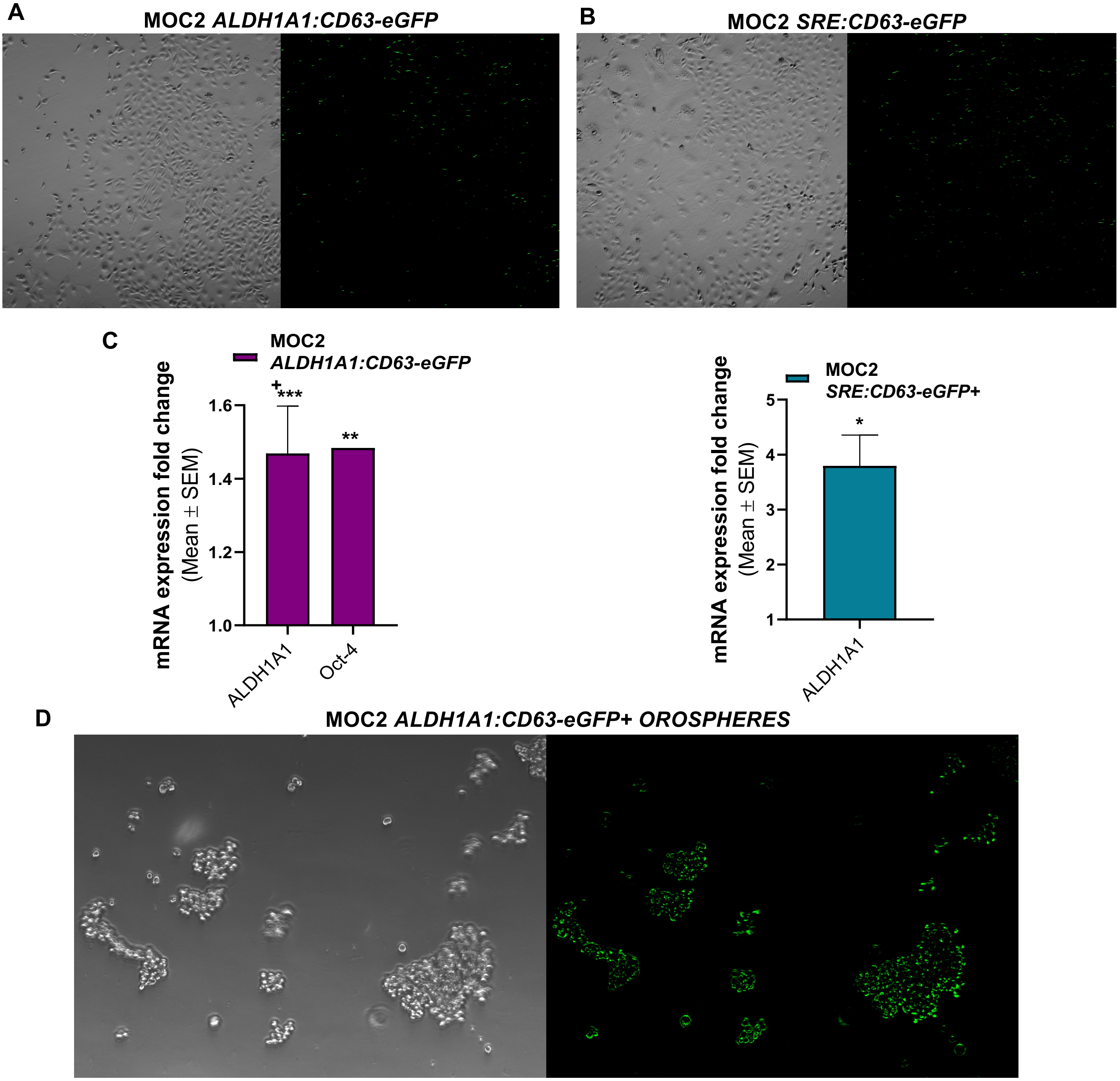
A
MOC2 ALDH1A1:CD63-eGFP
B
MOC2 SRE:CD63-eGFP
C
MOC2
ALDH1A1:CD63-eGFP
+
mRNA expression fold change
(Mean ± SEM)
1.0
1.2
1.4
1.6
***
**
ALDH1A1
Oct-4
MOC2
SRE:CD63-eGFP+
mRNA expression fold change
(Mean ± SEM)
1
2
3
4
5
*
ALDH1A1
D
MOC2 ALDH1A1:CD63-eGFP+ OROSPHERES

### Gating strategy

#### Tumor cells

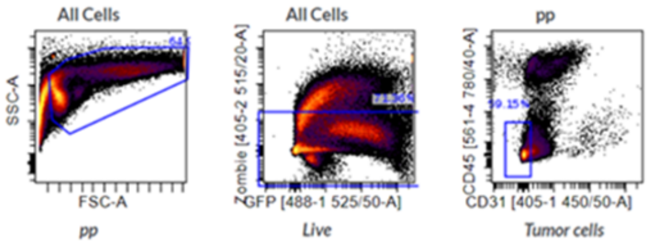


## EC

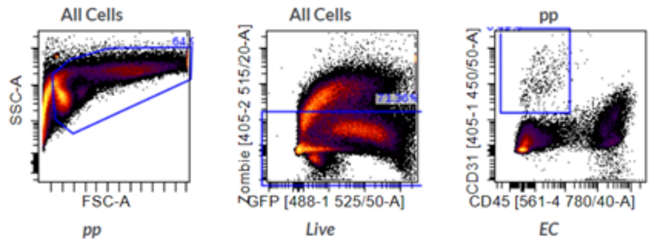


#### B cells

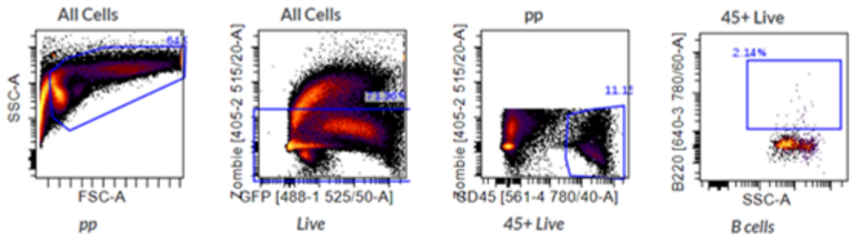


## DC

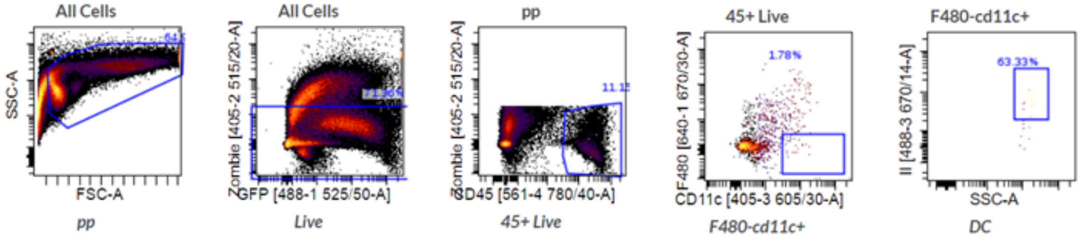


#### Neutrophils

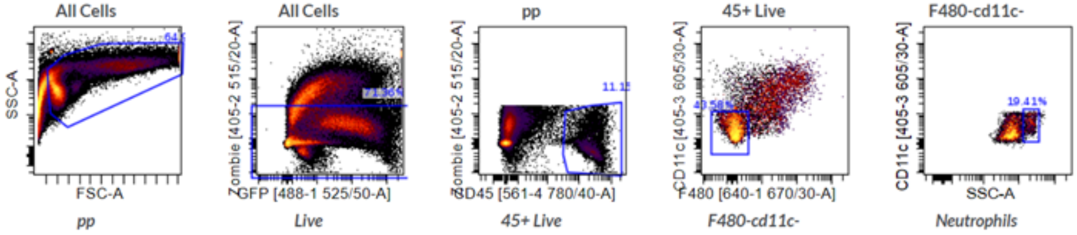

#### Inf Mo (11c+)

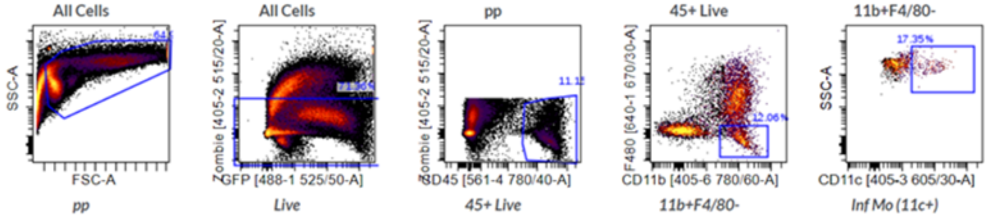


#### Res Mo (11c-)

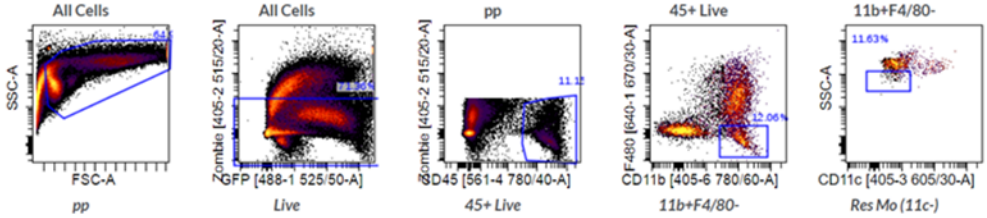


#### M1 Mac (II+)

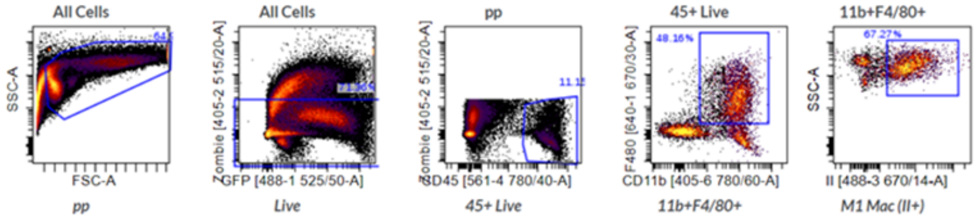


#### M2 Mac (II-)

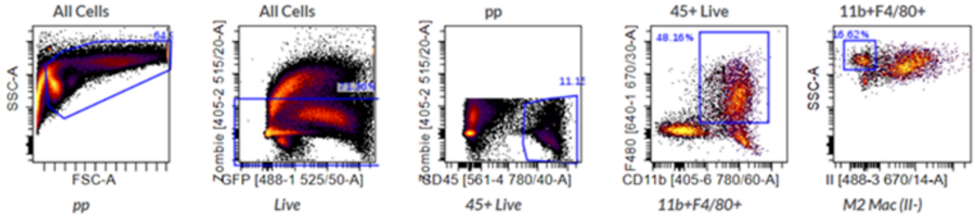


#### PD1+ T cells

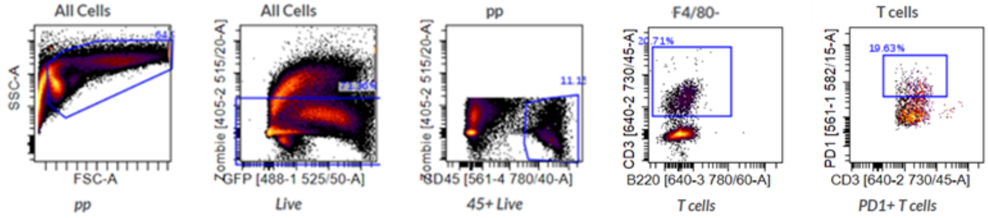


#### PD1- T cells

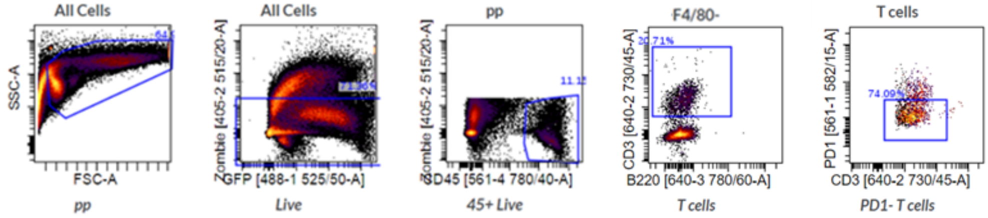

A

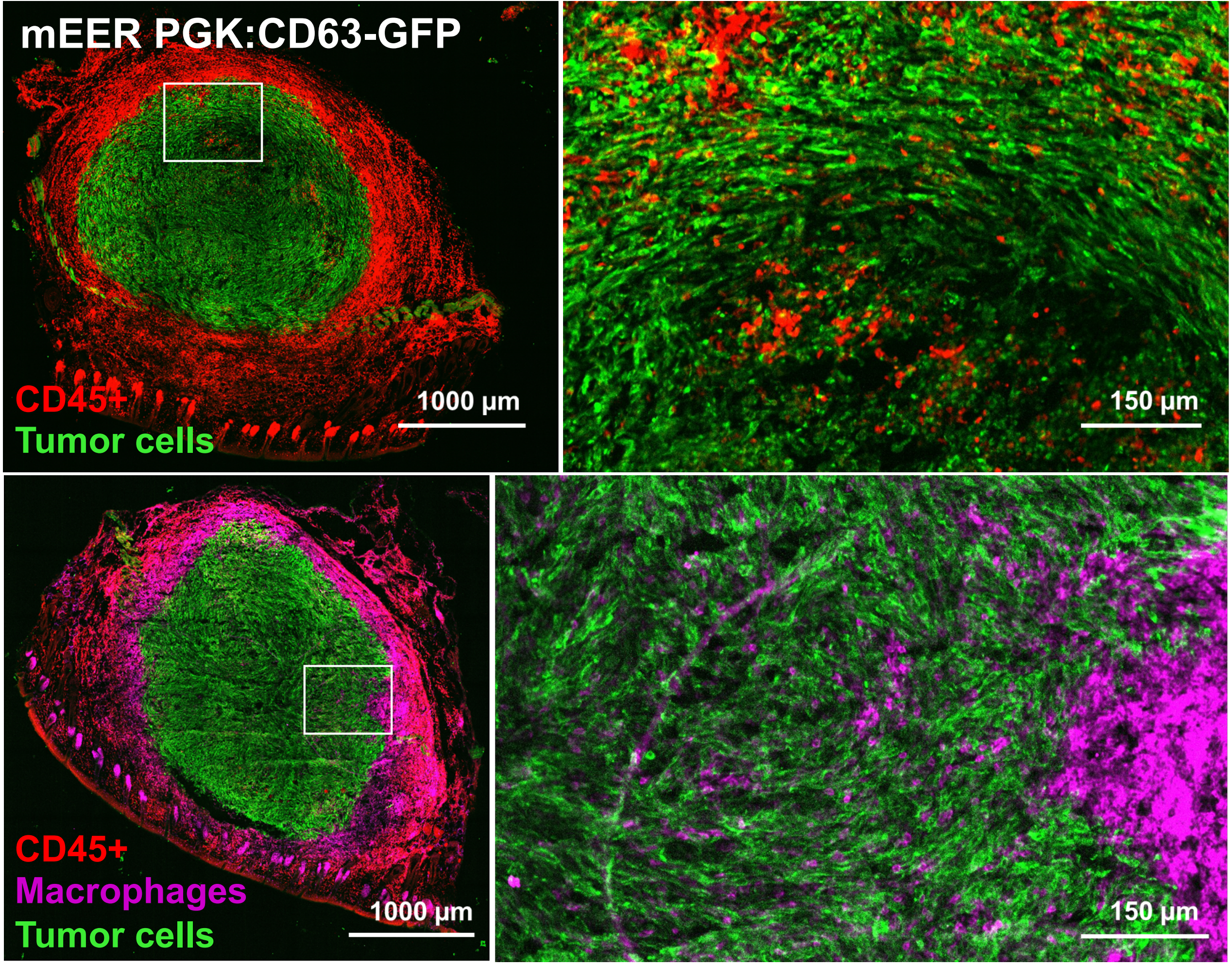
mEER PGK:CD63-GFP
CD45+
Tumor cells
1000 µm
150 µm
CD45+
Macrophages
Tumor cells
1000 µm
150 µm

B

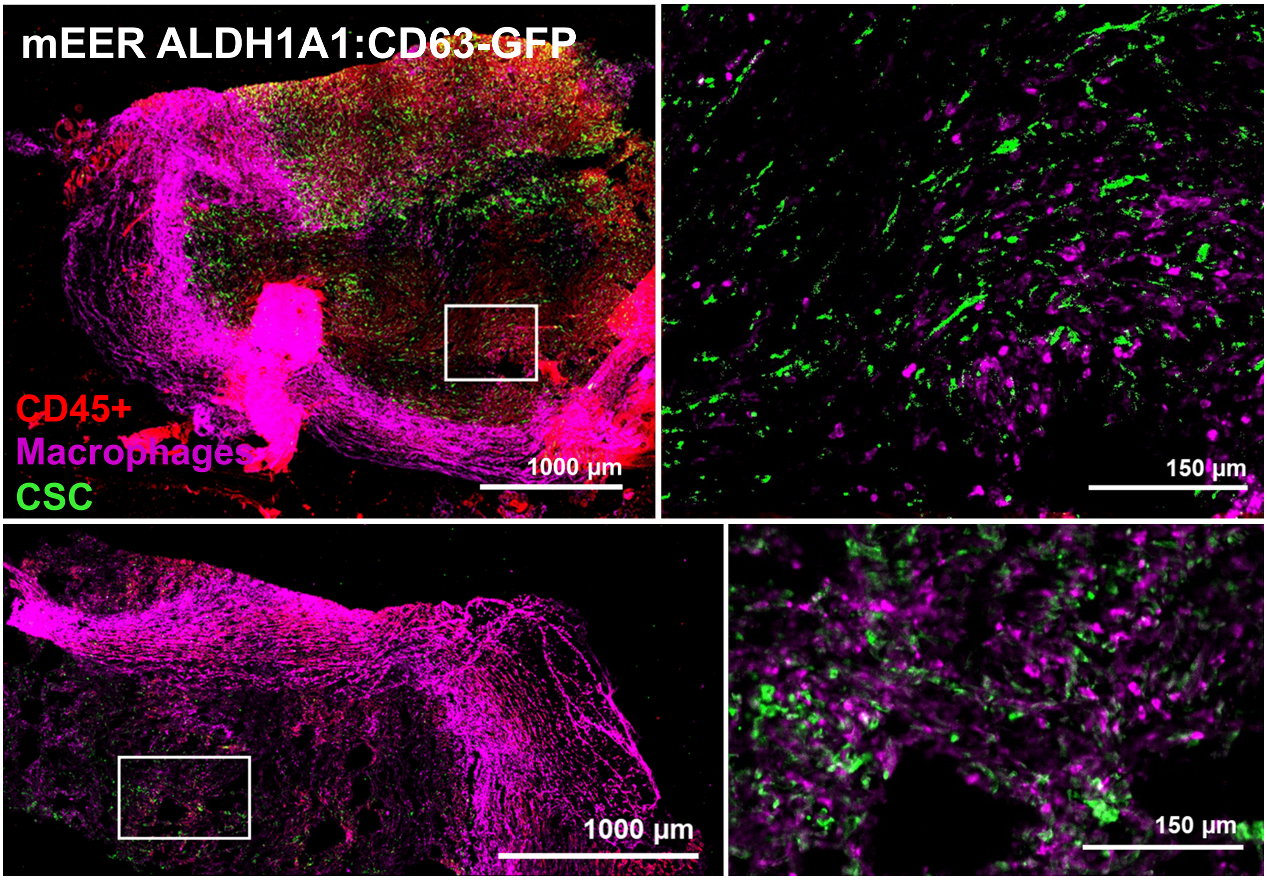
mEER ALDH1A1:CD63-GFP
CD45+
Macrophages
CSC
1000 µm
150 µm
1000 µm
150 µm
